# Transmission potential of Floridian *Aedes aegypti* mosquitoes for dengue virus serotype 4: Implications for estimating local dengue risk

**DOI:** 10.1101/2021.03.23.436716

**Authors:** Caroline J. Stephenson, Heather Coatsworth, Seokyoung Kang, John A. Lednicky, Rhoel R. Dinglasan

## Abstract

Dengue virus serotype 4 (DENV-4) circulated in *Aedes aegypti* in 2016 and 2017 in Florida in the absence of human index cases, compelling a full assessment of local mosquito vector competence and DENV-4 risk. To better understand DENV-4 transmission risk in Florida, we used an expanded suite of tests to measure and compare the vector competence of both an established colony of *Ae. aegypti* (Orlando strain [ORL]) and a field-derived colony from Collier County, Florida in 2018 (COL) for a Haitian DENV-4 human field isolate and a DENV-4 laboratory strain (Philippines H241). We immediately noted that ORL saliva-positivity was higher for the field versus laboratory DENV-4 strains. In a subsequent comparison with the recent COL mosquito colony we also observed significantly higher midgut susceptibility of COL and ORL for the Haitian DENV-4 field strain, and significantly higher saliva-positivity rate for COL, although overall saliva virus titers were similar between the two. These data point to a potential midgut infection barrier for the DENV-4 laboratory strain for both mosquito colonies and that the marked difference in transmission potential estimates hinge on virus-vector combinations. Our study highlights the importance of leveraging an expanded suite of testing methods with emphasis on utilizing local mosquito populations and field relevant dengue serotypes and strains to accurately estimate transmission risk in a given setting.

**Importance:** DENV-4 was found circulating in Florida (FL) *Ae. aegypti* mosquitoes in the absence of human index cases in the state (2016-2017). How DENV-4 was maintained locally is unclear, presenting a major gap in our understanding of DENV-4 public health risk. We determined the baseline arbovirus transmission potential of laboratory and field colonies of *Ae. aegypti* for both laboratory and field isolates of DENV-4. We observed high transmission potential of field populations of *Ae. aegypti* and evidence of higher vertical transmission of the DENV-4 field isolate, providing clues to the possible mechanism of undetected DENV-4 maintenance in the state. Our findings also move the field forward in the development of best practices for evaluating arbovirus vector competence, with evidence that transmission potential estimates vary depending on the mosquito-virus combinations. These data emphasize the poor suitability of lab-established virus strains and the high relevance of field-derived mosquito populations in estimating transmission risk.

## Introduction

Dengue viruses (DENVs) cause dengue fever, the most common mosquito-borne viral disease in humans. There are four dengue virus serotypes (DENV-1-4) that cause an estimated 400,000 infections globally [1]. DENVs are primarily transmitted by *Aedes aegypti* and human infections are commonly asymptomatic. Symptomatic patients can present with fever, rash, myalgia and/or arthralgia. Severe dengue (dengue hemorrhagic fever and shock syndrome) and death can develop from complications arising from immune enhancement following infection by a second serotype [2,3].

When *Ae. aegypti* bites a DENV-infected human, the virus travels with the blood meal to the mosquito midgut where it infects midgut epithelial cells [4,5]. This is followed by replication and dissemination into secondary tissues, culminating in infection of the mosquito’s salivary glands [6]. Physical barriers in the mosquito at each of these stages can influence DENV transmission success [4,7,8]. Mosquitoes inject saliva when taking a blood meal, but their bites are a route of DENV transmission only if virus is present in saliva. It is crucial to investigate a mosquito’s ability to become infected with DENV (their susceptibility), their ability to transmit DENV (their vector competence), and as a corollary, their transmission potential (determined as the presence of virus in saliva) to enhance our understanding of the biological drivers of arbovirus transmission in a local setting. *Ae. aegypti* arbovirus susceptibility and vector competence are not single measures within a mosquito species as they can vary across virus serotypes, virus strains and mosquito populations [9-13]. Identifying what these variations are within a mosquito population can help pinpoint virus features that confer higher infectivity, as virus strains with better fitness in mosquitoes have selective advantages for subsequent transmission to humans [14].

There have been sporadic outbreaks of dengue in Florida (FL, USA) since 2009, along with hundreds of imported travel cases from the Caribbean and Central America each year [15,16]. We reported the first instance of DENV-4 detection in recent history in Manatee County FL *Ae. aegypti*, and this was in the absence of any reported FL human index cases for this serotype, which could be pointing to virus maintenance in the mosquito population through vertical transmission [17]. The genome of this Manatee Co. DENV-4 strain has high nucleotide and amino acid sequence identity (>99%) with a strain isolated from Haiti in 2014, raising questions about the competence of FL *Ae. aegypti* to transmit this strain or similar strains. To date, few studies have examined DENV competence of *Ae. aegypti* in FL (1/32[∼3%] for DENV-1 BOLKW010) and the Caribbean (25/60 [42%] for DENV-4 H241) [12,18].

There are several gaps in the literature regarding DENV vector competence studies. First, the majority of published studies preferentially used a lab-adapted DENV-2, as reported in a review by Souza-Neto et al., 2019, biasing our understanding of mosquito competence for DENVs [9]. Second, there is a lack of experimentally proven “gold standards” for vector competence study methods for arboviruses, making comparisons difficult [9,19]. For example, it is common to provide a single infectious blood meal when ascertaining *Ae. aegypti* vector competence even though *Ae. aegypti* takes multiple blood meals per gonotrophic cycle. Recent work posits that current methods using only one blood feed may underestimate the transmission potential of arboviruses [20,21]. However, it remains unclear if subsequent blood feeding impacts transmission potential for all DENV serotypes equivalently, as only DENV-2 dissemination has been analyzed after a second blood feed [20]. Finally, the presence of insect-specific viruses (ISVs) such as Cell Fusing Agent virus (CFAV) in *Ae. aegypti* has been shown to influence vector competence [22]. Persistent CFAV infections have been detected in *Ae. aegypti* colonies and some arthropod cell lines [23,24], so estimates of vector competence could be confounded by ISVs such as CFAV if not accounted for. Given the lack of standardization and apparent limitations of current approaches, there is a clear need to move towards “best practice” to enable cross-regional comparisons of vector competence of *Ae. aegypti* populations.

Herein, we examined the susceptibility and transmission potential of a laboratory colony of *Ae. aegypti* from Orlando, FL (ORL) for a New World field isolate of DENV-4 from <u>H</u>aiti (DENV-4<u>H</u>) (GenBank accession MK514144.1) and an Old World prototype laboratory strain of DENV-4 (DENV-4<u>L</u>) (Philippines, H241, 1956), and assessed the utility of two methodological improvements in the rigorous measure of vector competence: the use of blood in collecting deposited virus during salivation and the requirement for repeated blood feeding on ORL transmission potential (virus in saliva). These studies were then extended towards the validation of the process with a recent, field-derived *Ae. aegypti* population from Collier County, FL. Taken together, we address the scarcity of knowledge about FL *Ae. aegypti* competence for DENV-4 and, in doing so, test several methodological improvements that can further augment and harmonize the rigor in determining DENV competence across the globe.

## Materials and Methods

### Mammalian cell culture and virus propagation

Vero E6 cells (African green monkey kidney epithelial cells, ATCC CRL-1586) were grown as monolayers at 37°C and 5% CO_2_ with aDMEM (advanced Dulbecco’s modified essential medium, Invitrogen, Carlsbad, CA), supplemented with 1% L-alanine-L-glutamine (GlutaMAX, Gibco, Gaithersburg, MD), 50 μg/mL penicillin, 50 μg/mL streptomycin, 100 μg/mL neomycin (PSN antibiotics, Invitrogen) and 0.25 µg/mL amphotericin B (Gibco) as well as 10% low-antibody, heat-inactivated (HI), gamma-irradiated fetal bovine serum (FBS, Hyclone, GE Healthcare Life Sciences, Pittsburgh, PA) (complete DMEM).

Experimental virus groups included a DENV-4 isolate from a symptomatic child in Haiti from 2015 (DENV-4H, strain *Homo sapiens*/Haiti-0075/2015, GenBank accession MK514144.1) and a DENV-4 Philippines/H241/1956 strain (DENV-4L, ATCC VR-1490). Virus stocks were propagated in Vero E6 cells with supplemented DMEM with 3% FBS (reduced DMEM) and collected on day 7 post-inoculation when approximately 50% of the cells were showing cytopathic effects. The stocks were clarified by centrifugation, frozen in a 10% trehalose solution, and cryopreserved in liquid nitrogen (Table S1). We did not observe any reductions in *in vitro* infectivity using cryopreserved virus stocks in infectious feeds shown by plaque assay replicates before and after freeze-thaw, unlike trends that have been described previously with ZIKV and DENV [25, 26].

### Mosquito rearing

*Aedes aegypti* Orlando strain (ORL) was collected from Orlando, Florida in 1952 and has since been reared at the United States Department of Agriculture’s Center for Medical, Agricultural and Veterinary Entomology (USDA-CMAVE) in Gainesville, Florida. *Ae. aegypti* Collier strain (COL) derived from pooled eggs collected by the Collier Mosquito Control District in September, 2018 from three different locales (Table S2). Mosquitoes were raised under standard laboratory conditions for mosquito rearing: 28°C with 80% relative humidity and a neutral photoperiod regimen (12 h light/12 h dark). Four to seven days post-eclosion, the mosquitoes were aspirated, cold-anesthetized and separated by sex so only females remained.

### Mosquito infection with DENV-4

Prior to blood feeding on day 0, mosquitoes were sugar-starved overnight. They were then artificially fed with 2:2:1, human hematocrit type O+ blood (hematocrit) (Lifesouth Community Blood Centers, Gainesville, FL): virus stock: heat inactivated (HI) human serum from warm glass feeders that were connected by tubing to a water bath set at 39°C. ORL and COL mosquitoes were blood-fed for one hour, then cold-anesthetized at 4°C for several minutes until immobile, and only blood-engorged mosquitoes were retained for further study. Mosquitoes were maintained as aforementioned for 7, 10, or 14 days post DENV blood feeding. All mosquito groups were given 10% sucrose and provided with a damp oviposition surface made from filter paper. Mosquito infections were performed in an Arthropod Containment Level 3 facility.

A separate 14-day ORL group of mosquitoes were fed an infected blood meal on day 0 [referred throughout as “1-feed”], while another group was similarly infected and then provided a non-infectious blood meal (1:1 hematocrit and HI human serum) 4 days after the infectious blood meal [referred throughout as “2-feeds”] (Figure 1, panel B).

**Figure 1.**
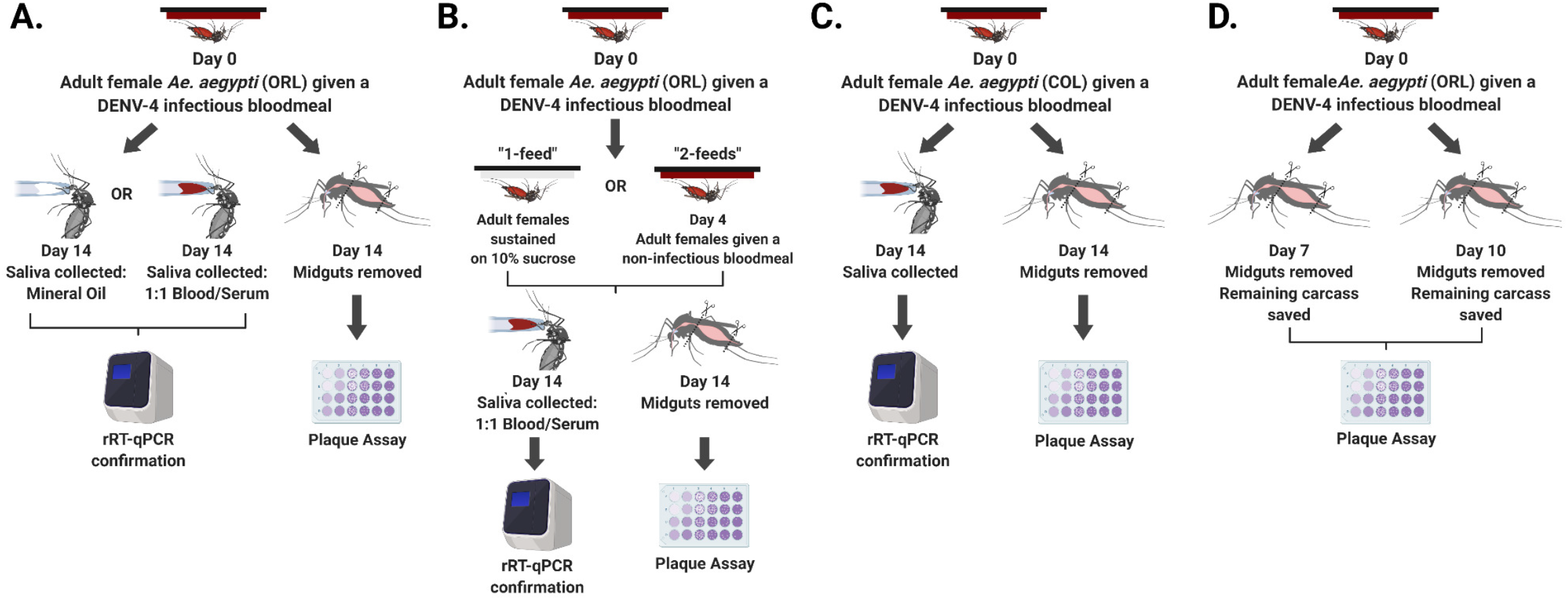
*Aedes aegypti* infection workflow. A) ORL day 14 saliva collection using mineral oil or blood, B) ORL day 14 collections comparing one and two blood feed groups, C) COL day 14 collections, and D) ORL day 7 and day 10 collections. Created with BioRender.com.

### Tissue preparations

Mosquito midguts were dissected 7-, 10-or 14-days post-infectious blood feeding. Individual dissected midguts were placed into 1.5 mL microcentrifuge tubes with 150 µL of reduced DMEM and approximately 1 gram of 0.5 mm sterile glass homogenization beads (Nextadvance, NY) [27]. The remainder of the mosquito body (containing the head, thorax, and abdomen, minus the midgut) was also similarly collected from the 7-and 10-day timepoints. All samples were immediately stored at −80°C until use (Figure 1, panel D).

### Mosquito saliva analysis

ORL and COL mosquitoes were sugar-starved overnight and then on 14-days post infectious blood-feed saliva was collected from mosquitoes in a manner similar to previously published methods [12,18,28]. In brief, ORL and COL mosquitoes were cold-anesthetized, and their wings and legs were removed with sterile forceps. Each mosquito was then fastened to a glass microscope slide with tape, and their proboscis was inserted into a graduated glass capillary tube (Drummond, Broomall, PA) filled with 3 µL of mineral oil (ORL only) or 1:1 hematocrit and HI human serum (ORL and COL) (Figure 1, panel A & C). Mosquitoes were then placed in a rearing chamber at conditions described above for forty-five minutes, or until they ingested approximately 2 µL of blood. Each proboscis was then removed from its capillary tube, and the blood from each capillary tube was aspirated into 1.5 mL microcentrifuge tubes with 200 µL of reduced DMEM. All samples were immediately stored at −80°C until use.

### Plaque assay

BHK-21 cells (Baby hamster kidney fibroblast cells, ATCC^®^ CCL-10^™^) were a kind gift from the Dimopulos laboratory (Johns Hopkins University) and were grown to confluency as aforementioned for Vero E6 cells and then seeded onto 24-well plates at a density of 5×10^4^/well and incubated at 37°C and 5% CO_2_ until newly confluent. Midgut or body samples from day 7, 10 or 14 from either the DENV-4H or DENV-4L experimental groups were homogenized using a Bullet Blender (Nextadvance, NY) adjusted to speed setting 8 for 3 minutes. Samples were then immediately centrifuged at an RCF of 2,400 for three minutes. Each sample was then serially diluted 10-fold in reduced DMEM and 100 µL of each dilution series was added to individual wells (Figure S1). The 24-well plates were rocked at room temperature for 15 minutes then incubated at 37°C and 5% CO_2_ for 45 minutes. Afterwards, 500 µL of reduced DMEM with 0.8% w/v methyl cellulose was added to each well and the plates were incubated for five days. On the fifth day, the spent media was removed from the 24-well plates and a 1:1 methanol/acetone solution with 1% w/v crystal violet added for one hour to fix and stain the cells. Plaques were counted manually, and the titer in plaque forming units/mL (PFU/mL) was determined.

### rRT-qPCR analyses

We sought to determine DENV-4 transmission potential by detecting virus genomic RNA (vRNA) in saliva samples via quantitative real-time RT-PCR (rRT-qPCR) from ORL and COL mosquitoes that were found to be midgut positive via day 14 plaque assays, under the assumption that a mosquito without a midgut infection could not have a disseminated infection or virus in its saliva. Total RNA was purified from 140 µL of each saliva sample using a QIAamp Viral RNA Mini Kit (Qiagen, Valencia, CA, USA), and eluted from the RNA binding columns using 80 µL of elution buffer. A final reaction volume of 25 µL (Superscript III, Invitrogen) containing 5 µL of purified RNA and the following dual target pan-DENV rRT-qPCR primers and probe (Table S3) was loaded as technical duplicates onto a BioRad CFX96 Touch Real-Time PCR Detection System. We estimated PFU equivalent (PFUe) for each DENV-positive saliva sample via regression analysis between PFU and C_q_ value of DENV-4H and DENV-4L stock viruses (explanation in the supplement). We then tested the eggs from ORL collected from filter paper that was provided as an oviposition surface throughout the 14-day period (completed in triplicate). Eggs were frozen at −80°C until assayed to ascertain the frequency of transovarial transmission (TOT) occurring during infection with these two virus strains between the 1-feed and 2-feeds groups. We randomly pooled 25 eggs in triplicate from each replicate, virus group and feeding group, resulting in 36 egg pools to test. Each egg pool was homogenized in 200 µL of 1X phosphate buffered saline (pH 7.4) with glass beads as described above, except with an extended time of six minutes total. We then extracted RNA and performed rRT-qPCR to determine DENV-4 positivity in ORL eggs.

### Presence of Cell Fusing Agent virus in mosquitoes

The mosquitoes used in these experiments were screened for CFAV via rtRT-qPCR since this virus can confound vector competence work (Table S3) [22].

### Statistical analyses

Reported results are pooled from three biological replicates for each experimental time point (workflow: Figure 1). We determined that pooling was appropriate via normality and homogeneity tests using GraphPad Prism version 8.0. All other statistical analyses were conducted using R statistical software [29], with the following libraries: FSA, rcompanion and multcompView [30,31,32]. Infection rates (IRs) (defined by the presence of viable virus in tissues as measured by plaque assay and reported dichotomously as “infected” or “not infected”) were analyzed via Fisher’s exact tests, while infection intensity (defined as the measured PFU or PFUe/mosquito between virus groups) was analyzed by non-parametric Kruskal Wallis and Dunn’s post-hoc tests for midgut and body titer data. Day 14 saliva titers were normally distributed (passed Shapiro-Wilk test) and were analyzed by a one-way ANOVA. All significance tests were performed at an α level of 0.05 and all infection intensity data were log transformed.

## Results

### Advantages of mosquito saliva collection into blood versus mineral oil

Harvesting of mosquito saliva in mineral oil is the most used collection method [9,18,28], yet many studies employing this method lack a positive control for salivation, and salivation into mineral oil is not a natural proxy for virus transmission during blood feeding. We postulated that compared to mineral oil, blood would be an improved collection medium given that it is a more realistic proxy for DENV transmission to humans and blood can be seen in the mosquito body post-feeding as a salivation control. We developed a robust protocol and compared this process and collection medium to the standard mineral oil methodology. There were no significant differences between saliva IRs for DENV-4H (*P* = 0.1429) between mineral oil and blood collection methods; however, for DENV-4L, since only one positive sample out of 39 for the mineral oil group was observed, statistical comparisons were not possible (Figure 2, panel A). DENV-4H infection intensities (virus titers in infected mosquitoes) were also statistically comparable between the two saliva collection methods (*P* = 0.07674) (Figure 2, panel B), but with a noticeably tighter confidence interval for DENV-4H collected into blood. The average DENV-4H saliva-positivity rate increased 4-fold from 7% to 30% following the switch to collecting saliva in blood and increased 9-fold from 2.5% to 23% for DENV-4L. Moreover, the rate of virus detection in saliva was more stable across replicates with collection into blood, for example DENV-4H rates were 6.6%, 0% and 16.6% for the three replicates of the mineral oil group and were 33.3%, 27.7% and 28.6% for the blood group. Therefore, collecting mosquito saliva into a capillary tube filled with blood results in a statistically comparable measure of transmission potential to the current field standard, yet yields a higher number of positive samples with stable rates across replicates. It also enabled the visualization of blood imbibement in the mosquito body immediately upon dissection.

**Figure 2.**
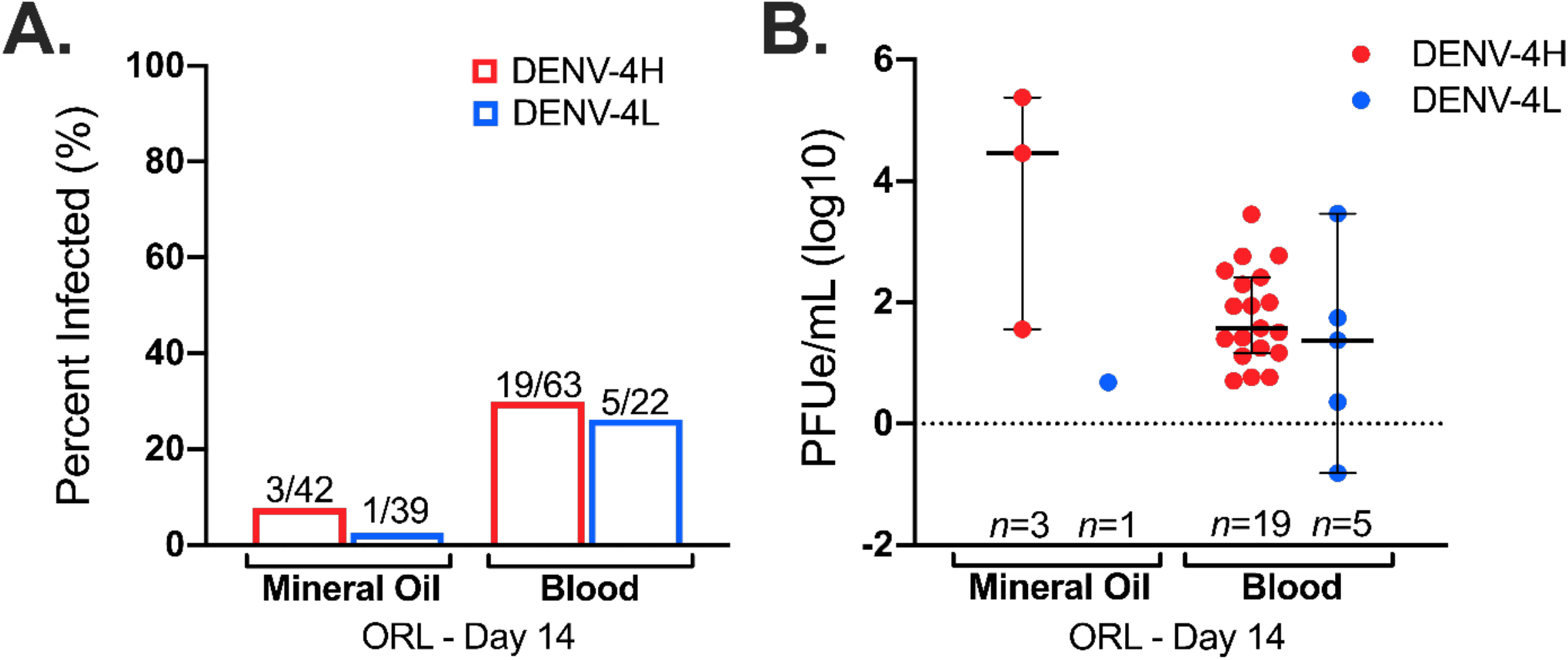
Collection of mosquito saliva into blood is statistically comparable to mineral oil but produced 4-fold (DENV-4H) and 9-fold (DENV-4L) higher positivity rates. A) day 14 infection rates and B) day 14 infection intensities of *Aedes aegypti* (ORL) saliva specimens with DENV-4 Haiti (H: red) and DENV-4 lab (L: blue), collected into capillaries filled with either mineral oil or blood. Dots represent individual mosquito samples, with median and 95% confidence intervals per group combined from three biological replicates.

### Additional bloodmeals do not significantly impact transmission potential or transovarial transmission of DENV-4

We determined if subsequent, non-infectious bloodmeals would affect estimates of the transmission potential of ORL for these two DENV-4 strains at 14 days post-infection. We leveraged the saliva collection method using blood as collection medium, to determine the 14 day transmission potential of DENV-4H and DENV-4L in ORL after only one infectious blood feed (1-feed) or following an additional non-infectious blood feed (2-feeds). DENV-4H midguts from both the 1-feed and 2-feeds groups had significantly higher IRs than both groups for DENV-4L (Figure 3, panel A). Neither pairwise comparisons between DENV-4H 1-feed versus 2-feeds IRs nor between DENV-4L 1-feed versus 2-feeds IRs were significantly different for midgut samples (Fishers exact test: *P* = 0.25 and *P* = 0.60, respectively) or saliva samples (*P* = 0.17 and *P* = 1).

**Figure 3.**
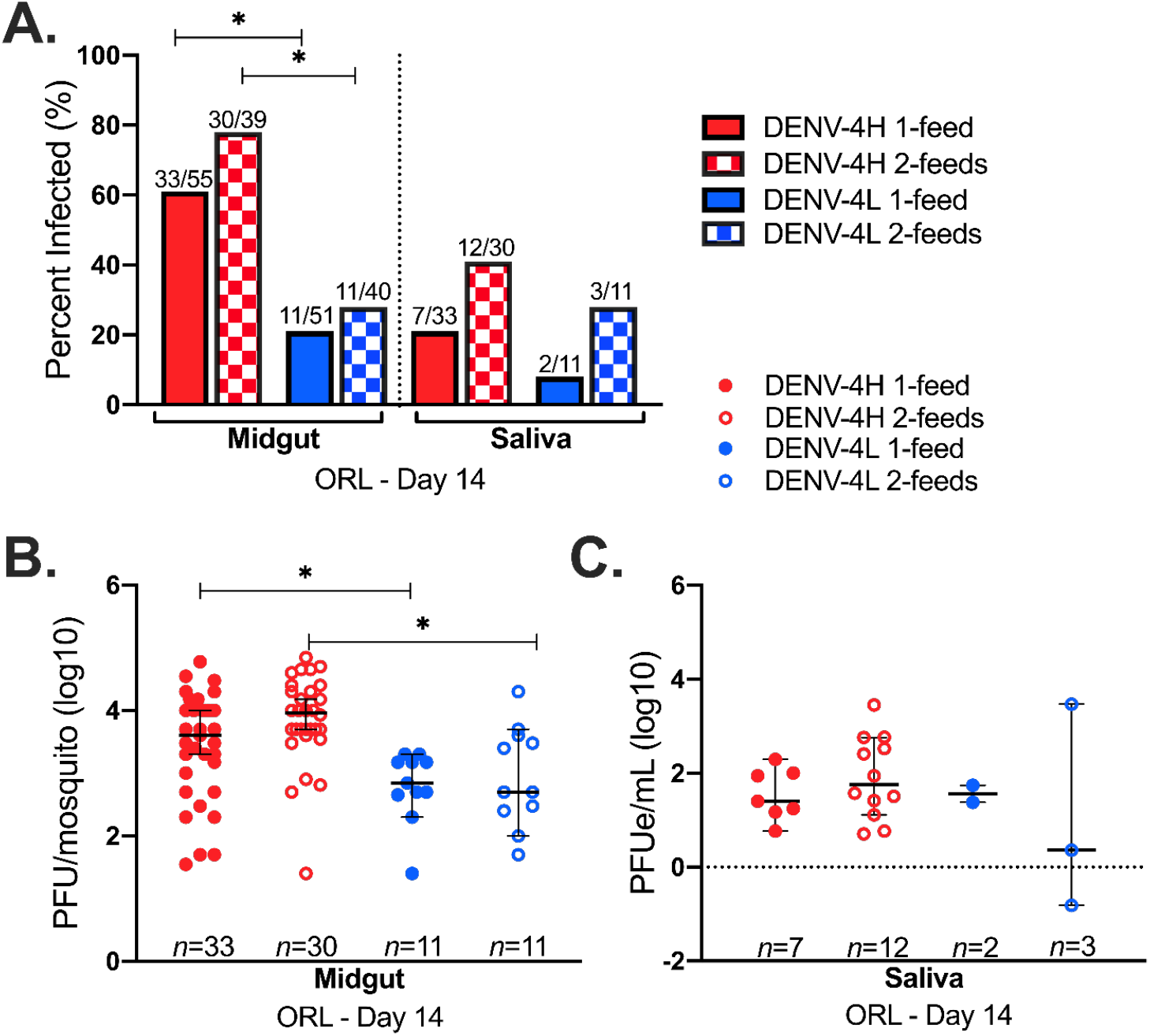
A successive non-infectious blood feed does not significantly impact transmission potential for DENV-4. *Aedes aegypti* (ORL) 14 day post-infectious blood feeding with DENV-4 Haiti (H: red) or DENV-4 lab (L: blue) (1 feed) and after a subsequent non-infectious blood feed four days later (2 feeds). A) Infection rates of midgut and saliva samples, B) midgut infection intensity, and C) saliva infection intensity. Dots represent individual mosquito samples, with median and 95% confidence intervals per group combined from three biological replicates. *P-value < 0.05 via Fisher’s exact test for infection rates and Kruskal-Wallis and Dunn’s post-hoc tests (midgut) or one-way ANOVA (saliva) for infection intensity.

Pairwise comparisons of infection intensities for day 14-collected midgut tissues between DENV-4H and DENV-4L 1-feed and 2-feeds groups revealed several significant differences (Kruskal-Wallis: *P* = 8.5×10^−5^) (Figure 3, panel B). Dunn’s post hoc test determined significant differences in measured midgut infection intensity between DENV-4H 1-feed versus DENV-4L 1-feed (*P* = 0.02), and between DENV-4H 2-feeds versus DENV-4L 2-feeds (*P* = 3.9×10^−3^). No significant differences between midgut titers for DENV-4L 1-feed versus 2-feeds (*P* = 0.10) nor between DENV-4H 1-feed versus 2-feeds (*P* = 0.054) were observed. The transmission potential of mosquitoes on day 14 with an established midgut infection was statistically comparable between DENV-4H and DENV-4L as well as between 1-feed versus 2-feeds groups (one-way ANOVA: *P* = 0.56) (Figure 3, panel C).

We then tested the eggs from ORL in each of these 1-feed and 2-feeds replicates to further understand if a successive blood feed would increase TOT success and if there would be a difference in TOT between the low passage DENV-4 field isolate versus the laboratory strain. One pool of ORL eggs tested positive for presence of DENV in duplicate PCR reactions from the DENV-4H 1-feed group. Two pools of ORL eggs tested positive for presence of DENV in duplicate PCR reactions from the DENV-4H 2-feeds group. In summary, ∼17% (3/18) of the pools tested positive for DENV-4H, while no egg pools (0/18) tested positive from any of the DENV-4L groups. Although the additional non-infectious bloodmeal does not appear to impact DENV-4 transmission potential or TOT, the overall trend in ORL infection and transmission potential for DENV-4H and DENV-4L appear to favor DENV-4H.

### Differential transmission potential of ORL and COL Ae. aegypti for DENV-4

Considering that ORL was colonized in 1952, it is unclear if vector competence data derived from the use of this colony line is representative of more current *Ae. aegypti* mosquito populations in FL. We compared transmission potential estimates of ORL to a recently colonized field population of *Ae. aegypti* from Collier Co. FL (COL). We observed significantly higher IRs for COL mosquito midguts infected with DENV-4H (64%) as compared to DENV-4L (21%) (Fishers exact test: *P =* 3.5×10^−5^), but not for saliva IRs (58% vs. 44%, respectively) (*P =* 0.21) (Figure 4, panel A). Midgut infection intensities were not significantly different between DENV-4H and DENV-4L (Kruskal-Wallis: *P =* 0.25) (Figure 4, panel B) nor were the DENV-4H or DENV-4L virus titer in saliva samples (*P* = 0.15) (Figure 4, panel C). In comparison to ORL, COL had significantly higher saliva-positivity rates for DENV-4H (*P* = 3.3×10^−5^) but all other comparisons between ORL and COL were similar.

**Figure 4.**
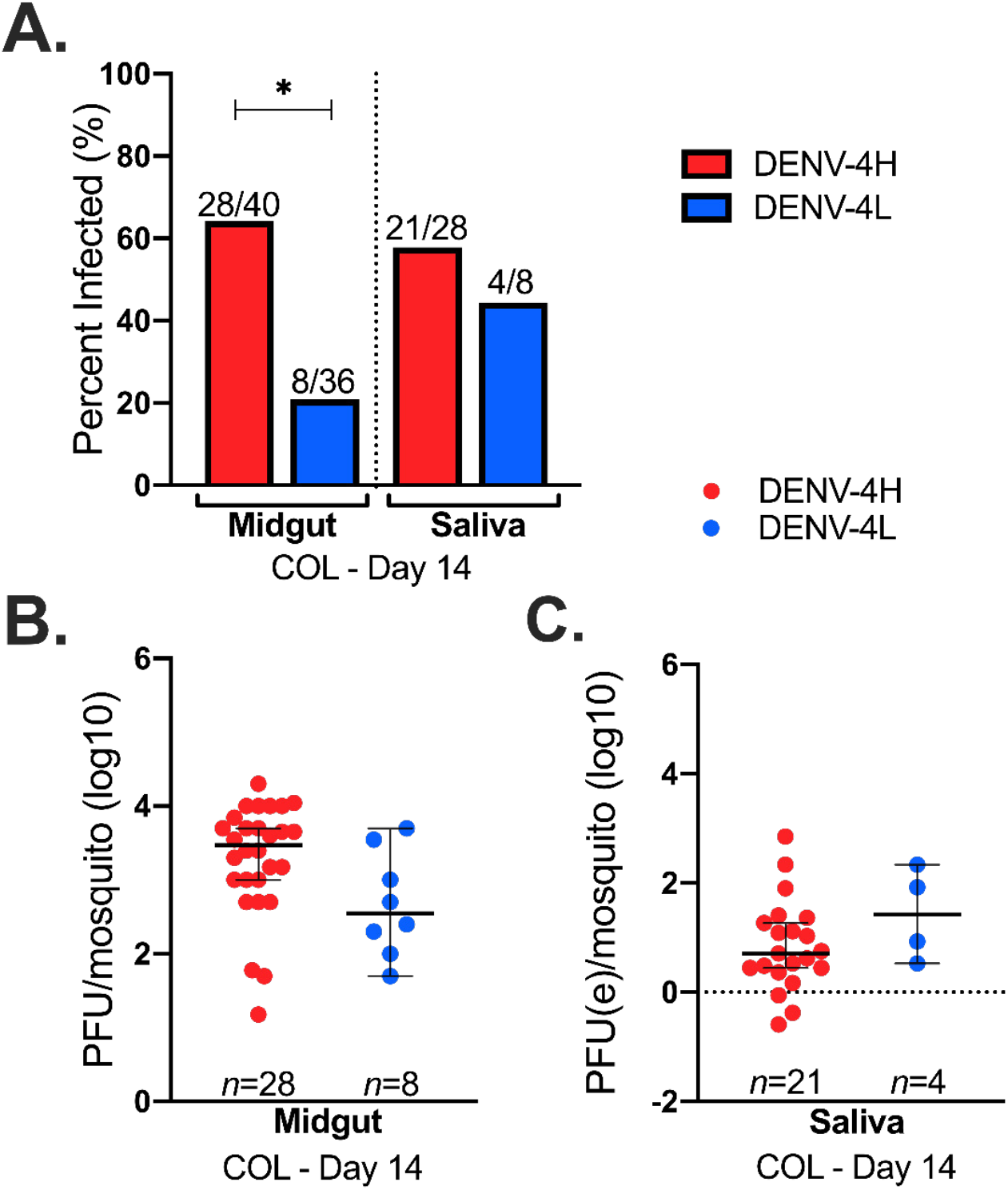
FL field mosquitoes have higher midgut susceptibility for DENV-4H than DENV-4L. Field-acquired *Aedes aegypti* (COL) 14 day post-infectious blood feeding with DENV-4 Haiti (H: red) or DENV-4 lab (L: blue). A) Infection rates of midgut and saliva samples, B) midgut infection intensity, and C) saliva infection intensity. Dots represent individual mosquito samples, with median and 95% confidence intervals per group combined from three biological replicates.*P-value < 0.05 via Fisher’s exact test for infection rates and Kruskal-Wallis and Dunn’s post-hoc tests (midgut) or one-way ANOVA (saliva) for infection intensity.

### Evidence of a midgut infection barrier for DENV-4L in ORL

Given the marked difference in susceptibility between DENV-4H and DENV-4L in both ORL and COL, we determined if these differences are apparent at earlier timepoints during infection and if there is differential dissemination of these viruses into the mosquito body. We focused on midgut versus body tissues on day 7 and day 10. Across all replicate studies, DENV-4H had significantly higher IRs in pairwise comparisons between midgut and body samples compared to DENV-4L (Figure 5, panel A).

**Figure 5.**
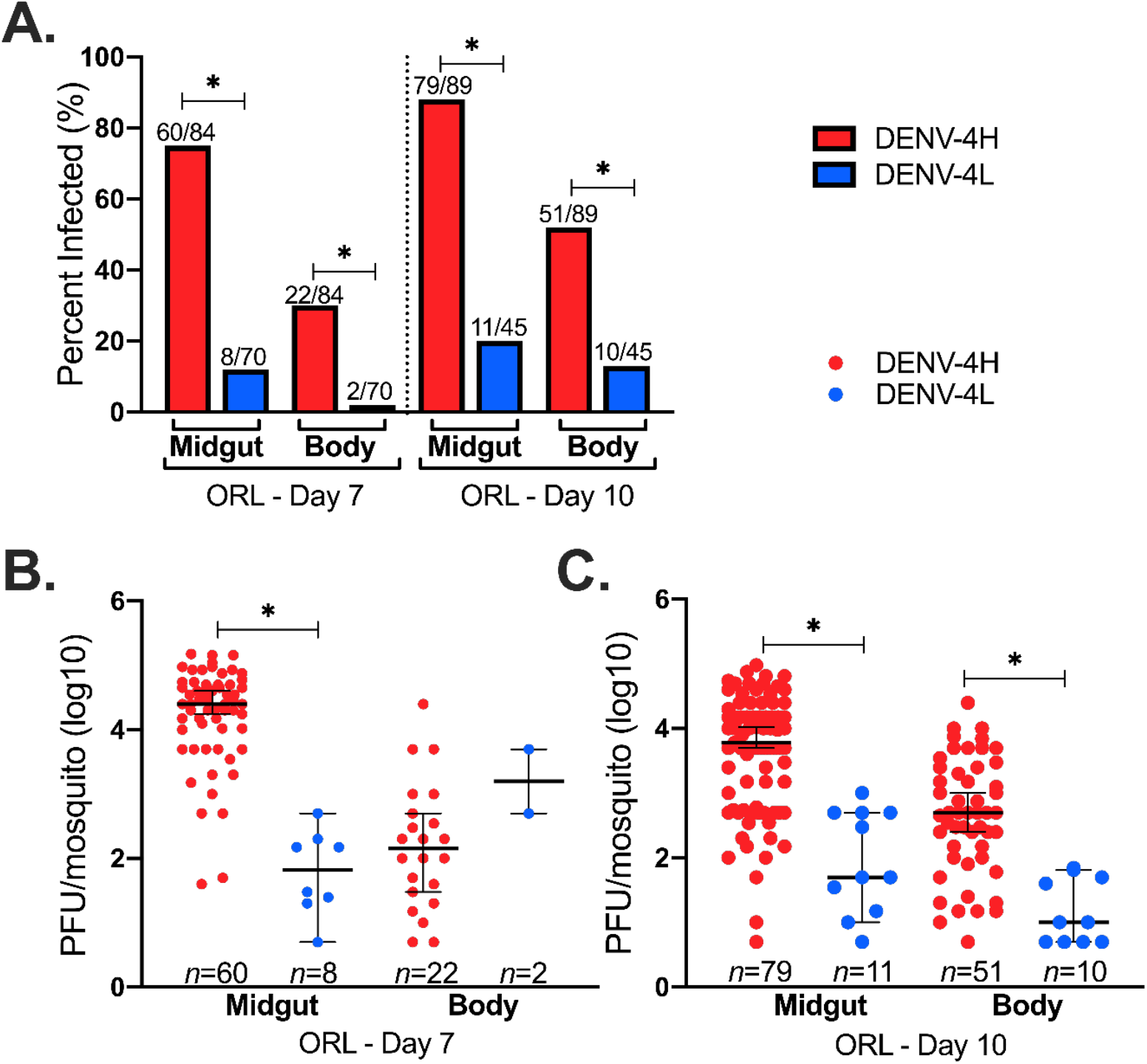
Significantly higher midgut susceptibility and dissemination for DENV-4H compared to DENV-4L points to a potential midgut barrier for the laboratory virus strain. *Aedes aegypti* (ORL) 7-and 10-day post-infectious blood feeding with DENV-4 Haiti (H: red) or DENV-4 lab (L: blue). A) Midgut and body infection rates, B) Day 7 midgut and body infection intensity, and C) day 10 midgut and body infection intensity. Dots represent individual mosquito samples, with median and 95% confidence intervals per group combined from three biological replicates. P-value < 0.05 via Fisher’s exact test for infection rates and Kruskal-Wallis and Dunn’s post-hoc tests for infection intensity.

The trends in infection intensity were similar to those observed for IRs. Virus titers were significantly higher for DENV-4H at all time points and in all tissues, except for day 7 body tissues where the difference was not statistically significant (Kruskal-Wallis: *p-value* (*P*) = 2.2×10^−16^) (Figure 5, panels B & C). These data show lower infection rates and slower dissemination for DENV-4L compared to DENV-4H.

## Discussion

We compared the susceptibility (infection of midguts), dissemination (infection of bodies), transovarial transmission (infection of eggs), and transmission potential (virus detection in saliva) of an isolate of DENV-4 from Haiti to the DENV-4 H241 prototype Asian lab strain in ORL *Ae. aegypti*. To understand the field comparability of these findings we then extended this study to a 2018 field-derived colony of *Ae. aegypti* from Collier Co. FL. It was crucial to determine the transmission potential of a FL field-derived *Ae. aegypti* population for DENV-4 to understand the risk posed to human health and gain insights into how this virus could become established in mosquito populations in the absence of reported human cases [17]. Importantly, we ruled out the contribution of CFAV infection, a potential confounder in vector competence studies, as neither of these tested mosquito strains had an active CFAV infection. To further enhance the rigor of the comparison, ORL mosquitoes were fed either one infectious blood meal only or one infectious blood meal plus one successive non-infectious blood meal several days later. There was higher DENV-4H susceptibility and dissemination in ORL compared to DENV-4L, and higher susceptibility for DENV-4H (versus DENV-4L) in COL. Transmission potential for DENV-4H in ORL was higher than DENV-4L but was not significantly different between groups receiving 1 vs. 2 bloodmeals. COL had a significantly higher transmission potential than ORL for DENV-4H but similar virus titer were measured in saliva between both mosquito strains across both virus strains. When transmission potential comparisons are made between the two virus groups for both ORL and COL mosquitoes with an established DENV-4 midgut infection on day 14 post-exposure, there were no significant differences for the likelihood of a mosquito having DENV-positive saliva. Therefore, the overall transmission potential between these two virus strains appear to be most impacted by midgut susceptibility and further work is needed to understand the specific mechanism that could be hindering DENV-4L infection and dissemination.

DENV-4H and DENV-4L share 93% nucleotide identity and 98% amino acid identity but the DENV-4H sequence has a notable 15-bp deletion in its 3’ untranslated region (similar to other DENV-4 genotype IIb strains that have spread globally) [17]. Minor genetic differences alone between these strains may have an impact on vector competence, as seen previously with CHIKV [33] and DENV [34]. Future work must ascertain if virus nucleotide changes are responsible for the differing phenotypes seen in our study, as these mutations may result in an enhanced midgut infection or escape barrier in ORL [4,34].

We determined that giving a successive blood meal during the EIP does not increase transmission potential, though it could still influence vectorial capacity by shortening the EIP [20], which is an area for further research. We observed no differences between the feeding groups in both the DENV-4H and DENV-4L comparisons between saliva samples collected on day 14 from ORL. This phenomenon could indeed vary across arboviruses, mosquito populations, and even between DENV serotypes, as this is the first reported successive blood feeding experiment to date for DENV-4 and mosquito susceptibility and competence varies considerably across serotypes and mosquito populations [9,12].

We developed our salivation assay protocols using published field standards as template [12,18,28,35]. Since we also tested blood as a medium for saliva collection, we were able to observe that mosquitoes had indeed salivated, as blood (a proxy for salivation as a natural process during blood ingestion) could be seen in the abdomen and confirmed upon dissection. Though, we were not able to determine quantity of saliva collected. We detected virus in saliva specimens via rRT-qPCR to avoid underestimation of saliva positivity rates due to inconsistent results comparing plaque assay and rRT-qPCR results for the same saliva samples, as reported by others previously [20].

We have identified considerable differences in ORL IRs of midgut and whole body samples, as well as in overall saliva positivity between field and lab strains of DENV-4. DENV-4H showed a higher occurrence of transovarial and horizontal transmission than DENV-4L. COL had similarly higher IRs for midgut and saliva samples for DENV-4H as compared to DENV-L. In both ORL and COL, it appears that a midgut barrier threshold is an initial hindrance to DENV-4L infection, but once an infection is established, the potential for transmission is similar to that of DENV-4H. Lastly, our study provides evidence that successive blood feeding during a 14-day EIP may not impact DENV-4 transmission equivalently for all vector-virus combinations. As such, we may have uncovered further serotype-specific nuanced responses related to transmission potential, which further underscores the importance of examining this parameter when estimating vector competence. Finally, transmission potential for COL was 58% for DENV-4H, on par with previous reports from the Caribbean [12], demonstrating that FL aegypti are efficient vectors as well as further emphasizing the importance of field arbovirus isolates in such studies. Pairing previous evidence of DENV-4 circulating in local *Ae. aegypti* populations [17] with our transmission potential data, we believe FL to be at-risk for local transmission of DENV-4, and that there are likely undetected pockets of DENV-4 maintenance throughout the state. Clearly, future studies measuring vector competence should consider investigating if similar trends also exist between other laboratory and fields strains of all DENV serotypes, especially if the goal is to accurately determine transmission risk of an arbovirus in a given setting.

## Acknowledgments

We thank the United States Department of Agriculture Center for Medical, Agricultural and Veterinary Entomology in Gainesville, Florida for providing *Ae. aegypti* (Orlando strain) and to Dr. Keira Lucas and the Collier Mosquito Control District for providing field-collected *Ae. aegypti* eggs for this work. We also thank Drs. Laurence Thirion and Remi Charrel from Unite des Virus Emergents, Aix Marseille University, France for providing primers and probes for the pan-dengue RT-PCR assay.

